# Toxicological study on methanol root bark extract of *Acacia sieberiana* (Fabaceae) in Wistar rats

**DOI:** 10.1101/2022.06.24.497563

**Authors:** Miriam Watafua, Jane I. Ejiofor, Aminu Musa, Mubarak Hussaini Ahmad

**Affiliations:** Department of Biochemistry, Faculty of Science, University of Maiduguri, Borno State, Nigeria; Department of Pharmacology and Therapeutics, Ahmadu Bello University, Zaria, Kaduna, Nigeria

**Keywords:** *Acacia sieberiana*, haematological indices, hepatic and renal biomarkers, histopathological investigations, sub-acute toxicity studies, weekly body weight

## Abstract

**Background:** The plant *Acacia sieberiana* belongs to the family Fabaceae. It has been used in ethnomedical practice to manage bleeding, rheumatism, pain, pyrexia, kidney diseases, gastrointestinal problems, parasitic and infectious diseases, hepatitis, cough, epilepsy, mouth ulcer and many more. Phytochemical compounds such as ellagic acid, quercetin, isoferulic acid, gallic acid, kaempferol, luteolin, apigenin, glucoside dihydroacacipetalin, acacipetalin and many others were isolated from Acacia sieberiana. Previous pharmacological investigations have reported that the plant has anticancer, antimicrobial, antidiarrhoeal and antitrypanosomal effects. Despite the therapeutic properties of this plant, no safety information is available in the literature. Hence, this work intends to investigate the sub-acute toxicity effects of *Acacia sieberiana* root bark extract (ASE). The phytochemical and oral median lethal dose (LD_50_) evaluations on the ASE were done in line with the standard protocols. The sub-acute toxic effects of the ASE (250, 750, and 1,500 mg/kg) were investigated following administration of the ASE daily for 28-consecutive days based on the Organization of Economic Cooperation and Development (OECD) 407 protocols in rats. The weekly body weights were monitored and the rats were euthanized on the 29^th^ day. The blood samples from the animals were obtained for biochemical and haematological determinations. The liver, kidney, lung and heart were removed for histological investigations.

**Results:** The ASE revealed triterpenes, tannins, saponins, cardiac glycosides, flavonoids, and alkaloids. The oral LD_50_ values was >5,000 mg/kg. The ASE remarkably (*p*<0.05) declined the body weight of the rats in consideration to the control categories. There was also a remarkable (*p*<0.05) elevation in ALP, urea and lymphocytes. The cardiac histology revealed no abnormalities. However, the liver produced dose-dependent hepatocellular necrosis and vacuolations. Besides, lymphocyte hyperplasia and glomerular necrosis were observed in the kidneys and alveolar congestion in the lungs.

**Conclusions:** The ASE is relatively non-toxic on acute administration. In contrast, it could pose slight hepatic and renal toxicity on sub-acute administration.

## 1.0 Background

The utilization of herbal plants as remedies has been practiced for years to obtain new therapeutic agents to manage various pathological disorders (Easmin et al., 2015). Even though there has been advancement in drug development procedures for use against ailments, still various diseases continue to affect the global population with significant death (Thomford et al., 2018). The herbal plants and their derivatives have been utilized in traditional practice by many people globally and serve as the main source of remedies against many diseases (Ahmad et al., 2021). It has been reported that about 80% of the global population utilizes herbal plants preparations as their significant cure to many pathological conditions (Kale et al., 2019). Besides, there has been an upsurge in the utilization of herbal preparations due to world attention towards phytotherapy, need for alternative medicine and the expensive cost of orthodox medications (Ouedraogo et al., 2012). Scientific investigations on herbal products have motivated the discovery of novel, potent and effective bioactive agents to treat diseases (Usman et al., 2021). About 30% of the active therapeutic compounds used clinically were obtained from herbal plants (Raskin et al., 2002).

There has been a perception that medicinal products obtained from natural sources including plants, are devoid of toxic consequences [8]. However, this general belief could be misleading as previous scientific studies have reported that many medicinal plants could produce various adverse effects including death (Bernstein et al., 2020; Chaachouay et al., 2020; Kharchoufa et al., 2018; Mounanga et al., 2015). Besides, there have been challenges to using herbal products such as defective standardization and dosage, insufficient identification and isolation, lack of clinical and toxicological profile of various medicinal plants (Okaiyeto & Oguntibeju, 2021; Thomford et al., 2018) such as *Acacia sieberiana*. As such, qualitative scientific evaluations on the harmful consequences of herbal plants and their derivatives used in traditional practice are required to develop new, effective and safe therapeutic agents (Ahmad et al., 2021; Ukwuani et al., 2012). Besides, the scientific information of herbal plants’ short and long term safety profiles is essential to document the safety concern of such plants for efficient therapeutic reasons (Reduan et al., 2020). Pharmacological agents must undergo preclinical screening with a battery of toxicological evaluations to give information on the potential safety of new compounds before human clinical trials (Denny & Stewart, 2017).

The plant *Acacia sieberiana* var Woodii (Fabaceae) is a tree of 3-25 m in height and 0.6-1.8 m in diameter (Ngaffo et al., 2020). The bark is rough, yellowish and peels off in rectangular, small and grey-brown scales with gummy exudates. The leaves are usually sparse and hairy and often bunched in pairs into small clusters from a common stalk, while the flowers are cream, white or pale yellow (Dawurung et al., 2012). It has dehiscent shiny brown fruits of about 1.3 cm thickness, 9-21 cm in length and 1.7-3.5 cm wide and which slowly splits open to release about 12 seeds (Dawurung et al., 2012). The plant grows in the savannah and appears with many botanical features throughout the Sahel and other African nations (Ameh & Eddy, 2014). It is drought and frost resistant and widely distributed in countries like Ethiopia, Benin, Chad, Gambia, Cameroon, Ghana, Kenya, Liberia, Zimbabwe, South Africa, Mozambique, Senegal, Mali, Mauritania, Namibia, Sierra Leone, Swaziland, Sudan, Nigeria, Portugal, Tanzania, Uganda, Zambia, Togo and India (Ameh & Eddy, 2014). In Nigeria, it is grown extensively as an economic tree in the Northern regions especially in Yobe, Jigawa and Sokoto States (Ameh & Eddy, 2014). The common names of the plant are umbrella thorn/white thorn/paperback thorn/flat-topped thorn or paperback (English); *Farar kaya* (Hausa); *Aluki* or *Sie* (Yoruba); *siyi* (Igbo); *Daneji* (Fulani); *Umkhaya* (Zulu) (Ameh & Eddy, 2014; Salisu et al., 2014).

In African traditional practices, the plant gained popularity in the treatment of many ailments. The roots decoction is utilized traditionally to treat bilharzia, bleeding, gonorrhoea, syphilis, rheumatism, kidney diseases, ophthalmia, stomach-aches, acne, tapeworms, urethral problems, and oedema (Ngaffo et al., 2020). The root decoction is also used in Nigeria to treat hepatitis (Ohemu et al., 2014). The powdered bark is used against fever in paediatric patients. The *Acacia sieberiana* is used in Nigeria to treat diarrhoea (Offiah et al., 2011). The pods serve as an emollient and as astringent. *Acacia sieberiana* has been used in traditional practice to manage skin eruptions, gastritis, cough, ringworm, leprosy, convulsion, dysentery and oral ulcer (Obidah et al., 2009). The bark and stem of the *Acacia sieberiana* are utilised to treat fever, jaundice, impotence, erectile dysfunction, syphilis, haemorrhoids, schistosomiasis, and enhance milk production after child delivery (Dawurung et al., 2012).

Previous phytochemical investigation on the *Acacia sieberiana* leaves leads to the isolation of gallic acid, kaempferol, ellagic acid, quercetin, isoferulic acid, kaempferol 3-α-_L_-arabinoside and quercetin 3-O-β-_D_-glucoside (Abdelhady, 2013). Other compounds including luteolin-7-O-rutinoside, chrysoeriol-7-O-rutinoside, apigenin-7-O-β-_D_-glucopyranoside, chrysoeriol-7-O-β-_D_-glucopyranoside, luteolin, luteolin-3′,4′-dimethoxylether-7-O-β-_D_-glucoside and sitosterol-3-O-β-_D_-glucoside, were isolated from the leaves of the plant (Ngaffo et al., 2020). The stem and leaves of *Acacia sieberiana* contain a glucoside dihydroacacipetalin and acacipetalin (Seigler et al., 1975).

The leaves extract of *Acacia sieberiana* elicited anticancer activity (Ngaffo et al., 2020). The stem, roots, bark and leaves of *Acacia sieberiana* were reported to have antibacterial (Kirabo et al., 2018) and antidiarrhoeal actions [18,27]. Besides, the stem bark extract of Acacia sieberiana exerted antitrypanosomal activity (Ogbole et al., 2021). The plant has been ingested as a nutritional source of protein in North-Western Nigeria (Salisu et al., 2014).

Despite the ethnomedicinal indications of *Acacia sieberiana* in traditional practice and therapeutic potentials against various diseases, there is unavailability of safety data on its sub-chronic administration. Hence, the current study investigated the sub-acute toxicological profile of methanol roots bark extract from *Acacia sieberiana* to bring out information on its safety for traditional utilization and activate more investigations to develop new, efficacious and safe bioactive agents against various ailments.

## 2.0 Methods

### 2.1 Collection and identification of the plant

The *Acacia sieberiana* plant was sourced from Samaru, Zaria, Kaduna State of Nigeria, identified (voucher specimen number: 16136) by Mallam Namadi Sanusi at the Department of Biological Sciences, Ahmadu Bello University (ABU), Zaria, Nigeria.

### 2.2 Animals

Adult Wistar rats (males and females) weighing between 120 g to 200g were obtained from the Animal House of the Pharmacology and Therapeutics Department, ABU, Zaria, Nigeria. They were kept in sufficiently-ventilated polypropylene cages, provided with sufficient animal diet (Vital feed, Jos, Nigeria) with water provision *ad libitum*. The rats were kept at appropriate laboratory situations (temperature 22 ± 3 °C, relative humidity of 30-70%) for fourteen days to acclimatize to the laboratory environment before the experimental methods. All the experimental procedures complied with the ABU Ethical Committee on Animal Use and Care Research Policy (ABUCAUC) and ARRIVE (Animal Research: Reporting of *In Vivo* Experiments) guidelines. The ethical approval number for the experiment is ABUCAUC/2016/049. Immediately following the experiment, all the rats were anaesthetized with chloroform, quickly euthanized by cervical dislocation and buried as per the University’s guideline for appropriate disposal of experimental animals remain.

### 2.3 Plant preparation and extraction

The collected fresh root barks of *Acacia sieberiana* were washed with clean water to get rid of contaminants, then air-dried in a shaded condition to a uniform weight and size-reduced into a coarse powder with the aid of mortar and pestle. The powdered sample (2,500 g) was soaked in 10 litres of 70%v/_v_ methanol in a conical flask for 72-hours with frequent shaking to stir. The Whatman filter paper (No. 1) was used to filter the mixture, and the resultant filtrate was concentrated to a constant weight on a water bath maintained at 50□C. The obtained extract was kept packaged and labelled as *Acacia sieberiana* extract (ASE). A fresh solution of the ASE was prepared on each day of the experiment with distilled water.

Then the extractive value of the ASE was evaluated as follows:

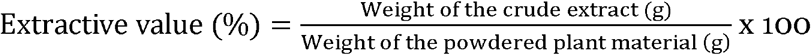

### 2.4 Phytochemical evaluation

Preliminary phytochemical evaluation on ASE was done to identify the phytochemical constituents using an appropriate procedure (Sofowora, 1993).

### 2.5 Acute toxicity evaluation

The acute toxicity determination on the methanol root bark extract of *Acacia sieberiana* was evaluated in rats after oral administration as per Lorke’s method (Lorke, 1983). The acute toxicity effect of the ASE was conducted in two phases with 12 rats. In the 1^st^ phase, 9 rats were categorized into 3 different groups (n=3 rats per group) and administered with the ASE orally at the doses of 10, 100 and 1000 mg/kg. The rats were observed for 24-hours for signs of harmful effects or loss of life. With the absence of death in the 1^st^ phase, the 2^nd^ phase of the test was conducted, with 3 rats categorized into 3 groups (n=1 rat per group) and administered with the higher doses of the ASE orally (1600, 2900 and 5,000 mg/kg) in a respective manner. The rats were eventually observed for 24-hours for possible signs of harmful effects and loss of lives. The oral median lethal dose (LD_50_) was estimated by taking the geometric mean of the highest non-lethal and the lowest lethal doses as follows:

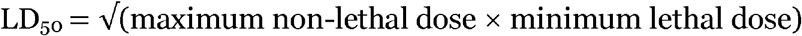

### 2.6 Sub-acute toxicity study

The sub-acute toxic effects of the ASE was done as per the Organization of Economic Co-operation and Development (OECD) 407 procedure for testing compounds (OECD, 2008). Forty (40) rats were categorised into four classes of 10 rats per group (5 males and 5 females). The 1^st^ category is the control received distilled water (1ml/kg) orally, whereas the 2^nd^, 3^rd^ and 4^th^ classes received 250, 750, and 1,500 mg/kg of the ASE, respectively, once daily for 28-consecutive days using an orogastric cannula. The rats’ body weights were monitored weekly, and there was observation for possible signs and symptoms of harmful actions and death. On the 29^th^ day, the animals were anaesthetized with chloroform after being deprived of food overnight with adequate water. The blood samples were obtained via cardiac puncture into ethylenediaminetetraacetic acid (EDTA)-containing and plain tubes for blood and biochemical investigation, respectively. The animals were euthanized immediately by cervical dislocation and buried deeply as per the University’s guide for appropriate disposal of experimental animal remains.

#### 2.6.1 Biochemical analysis

Blood for the biochemical analysis was kept at room temperature for 1-hour to clot and centrifuged at 3,000 revolutions per minute (rpm) for 10 minutes. The resultant serum was used to investigate the levels of liver parameters such as aspartate aminotransferase (AST), alanine aminotransferase (ALT), alkaline phosphatase (ALP), total protein (TP), albumin (Alb), total and direct bilirubin as well as for renal indices including creatinine, urea and electrolytes (potassium, chloride, sodium and bicarbonate ions).

#### 2.6.2 Haematological analysis

Blood parameters such as red blood cells (RBC), haemoglobin (Hb), platelet, white blood cells (WBC), packed cell volume (PCV) and differentials such as neutrophils, lymphocytes, monocytes, eosinophils, basophils were analyzed using an automated haematology analyzer.

#### 2.6.3 Histopathological studies

The excised organs (kidney, liver, heart and lung) were fixed in 10% formalin solution for histological examination. The sectioning of the organs (4-5μm) was done using a microtome to cut very fine sections of the embedded tissues, which were then floated out on a water bath and placed on microscope slides. The slides were dried on a hot plate to get rid of moisture for the tissue to adhere to the slide. De-waxing was done by using a solvent to remove the wax from the slide before staining. The tissue on the slide was then stained with haematoxylin and eosin (H&E) and covered with a glass to make the preparation permanent. The tissue slides were then viewed at a magnification of ×250 and photomicrographs of the tissues were obtained.

#### 2.7 Data analysis

The values generated were represented as mean ± standard error of the mean (SEM) in tables. One way analysis of variance (ANOVA) was used to analyze the biochemical and haematological parameters; whereas; split-plot ANOVA was used to analyze the weekly body weights followed by Dunnet’s multiple comparison *post hoc* test. The p≤0.05 was taken as a level of statistical significance.

## 3.0 Results

### 3.1 Extractive yield

A sticky dark-brown solid residue weighing 115.78g (4.63%^w^/_w_) with a mild sweet smell was obtained from 2,500g crude root bark of the *Acacia sieberiana* powdered sample.

### 3.2 Phytochemical constituents

Preliminary phytochemical screening of the *Acacia sieberiana* root bark extract showed cardiac glycosides, triterpenes, tannins, flavonoids, saponins and alkaloids, while anthraquinones and steroids were absent.

### 3.3 The oral median lethal dose (LD_50_)

Acute oral administration of the ASE showed no death at doses up to 5,000mg/kg and thus, the oral LD_50_ of the extract was estimated to be ≥5,000 mg/kg body weight, and there were no signs of abnormal behaviour.

### 3.4 Weekly body weights

There was an increase in the weight of the animals in each group over the four weeks. However, the animals treated with the extract showed lesser weight gain relative to the control group. In week 1, the mean weight of all the extract-treated groups was almost similar. However, the ASE at all the doses showed remarkable (*p*<0.05) weight reduction in the subsequent weeks (2, 3 and 4) in relation to the control group. The actions of the ASE on the weekly body weight of the animals following 28-days repeated oral administration are presented in Table 1.

**Table 1:**
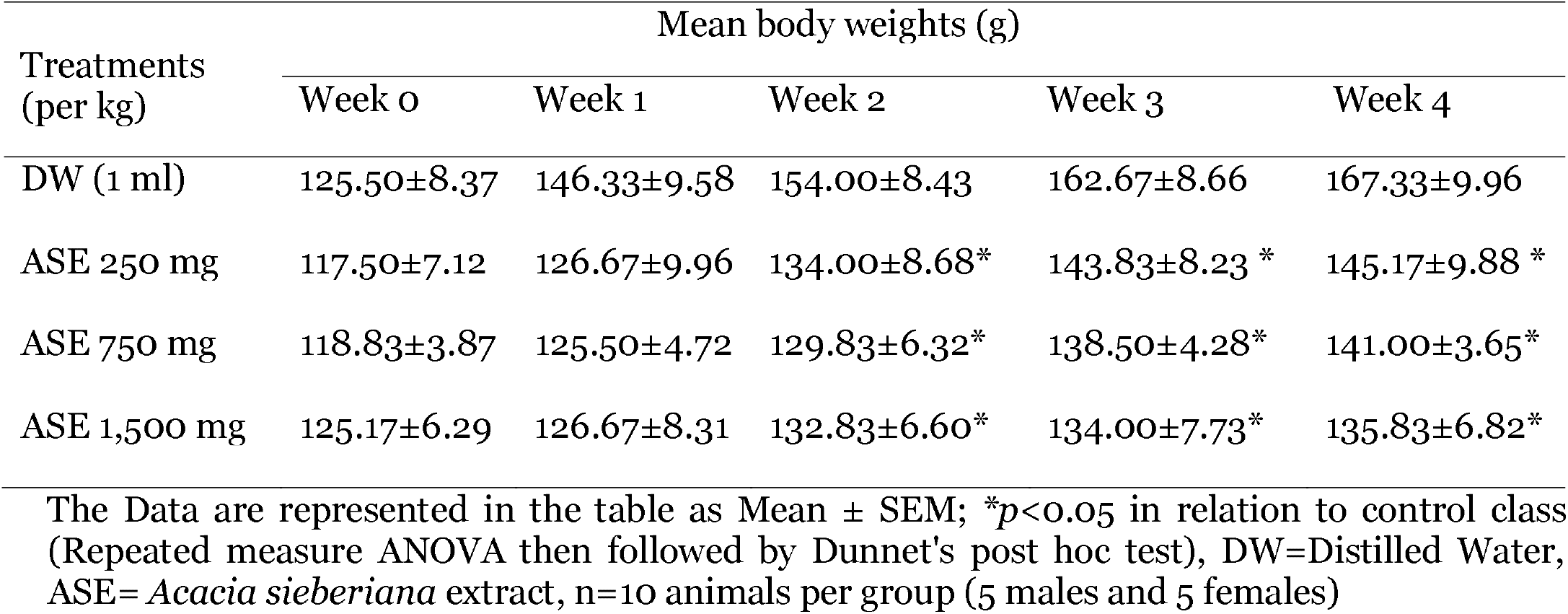
Weekly body weights.

### 3.5 Liver parameters

The ALT, AST, total protein, albumin, and bilirubin (direct and total) were within normal physiological values relative to the control group. On the contrary, a remarkable (*p*<0.05) and dose-dependent elevation in serum ALP level occurred relative to the control group. The effects of the ASE on the hepatic parameters following 28-days of repeated oral administration are shown in Table 2.

**Table 2:** Liver parameters.

| Liver parameters | Treatment groups (per kg) |  |  |  |
| --- | --- | --- | --- | --- |
|  | D/W (1 ml) | ASE 250 mg | ASE 750 mg | ASE 1,500 mg |
| ALT (IU/L) | 6.00 ± 0.68 | 7.17 ± 0.80 | 8.00 ± 1.24 | 7.33 ± 0.67 |
| AST (IU/L) | 12.67± 0.92 | 12.00± 0.58 | 11.00± 1.67 | 10.33± 1.28 |
| ALP (IU/L) | 26.76± 3.38 | 43.78±4.32* | 45.40±3.24* | 50.21±4.03* |
| Total Protein (g/dL) | 6.54±0.24 | 6.69±0.32 | 6.44±0.30 | 6.93±0.34 |
| Albumin (g/dL) | 3.14±0.07 | 3.17±0.08 | 3.08±0.06 | 3.03±0.09 |
| Direct Bilirubin (mmol/L) | 0.34±0.06 | 0.26±0.03 | 0.24±0.04 | 0.30±0.02 |
| Total Bilirubin (mmol/L) | 0.56±0.08 | 0.52±0.06 | 0.42±0.07 | 0.50±0.06 |

The data were represented as Mean ± SEM; \* $p < 0.05$ compared to control group (One way-ANOVA then followed by Dunnet's post hoc test), DW=distilled water, ASE= *Acacia sieberiana* extract, AST=Aspartate aminotransferase, ALT=Alanine aminotransferase, ALP=Alkaline phosphatase, I.U=International Unit, ASE=*Acacia sieberiana* extract, n=10 animals per group (5 males and 5 females)

### 3.6 Kidney parameters

This result showed a dose-related and remarkable (*p*<0.05) increase in serum urea at 1,500 mg/kg as related to the control group. The creatinine and all the electrolytes were within the normal limit. The effects of the ASE on the renal parameters following 28-days of repeated oral administration are presented in Table 3.

**Table 3:** Kidney parameters.

| Kidney biomarkers | Treatment groups (per kg) |  |  |  |
| --- | --- | --- | --- | --- |
|  | D/W (1 ml) | ASE 250 mg | ASE 750 mg | ASE 1,500 mg |
| Urea (mg/dL) | 29.38± 1.22 | 29.80 ± 0.89 | 31.50 ± 2.70 | 34.23 ± 1.82* |
| Creatinine (mEq/L) | 0.97±0.05 | 1.03±0.11 | 0.97±0.12 | 1.24±0.10 |
| K<sup>+</sup> (mmol/L) | 23.49± 2.10 | 20.66± 1.83 | 22.78± 1.36 | 24.87± 2.27 |
| Na<sup>+</sup> (mmol/L) | 225.32± 7.44 | 225.26±2.99 | 223.08±6.91 | 224.83±8.24 |
| Cl<sup>-</sup> (mg/dL) | 96.67±5.39 | 98.67±5.26 | 87.83±3.02 | 89.40±1.50 |
| HCO<sub>3</sub><sup>-</sup> (mg/dL) | 22.17±1.33 | 23.00±1.77 | 22.17±1.35 | 21.00±1.92 |

The data were represented as Mean ± SEM; \* $p < 0.05$ compared to control group (One-way ANOVA then subsequently by Dunnet's post hoc test), DW=Distilled water, ASE= *Acacia* *sieberiana* extract, K<sup>+</sup>=potassium ion, Na<sup>+</sup>=sodium ion, Cl<sup>-</sup>=chloride ion, HCO<sub>3</sub><sup>-</sup>=bicarbonate ion, n=10 animals per group (5 males and 5 females)

### 3.7 Haematological parameters

The extract has no effects on PCV, Hb, WBC, RBC, platelet, neutrophils, eosinophils and monocytes as related to the control group. However, the levels of lymphocytes elevated at the dose of 1,500 mg/kg in relation to the control group. The effects of the ASE on the blood parameters following 28-days of repeated oral administration are presented in Table 4.

**Table 4:**
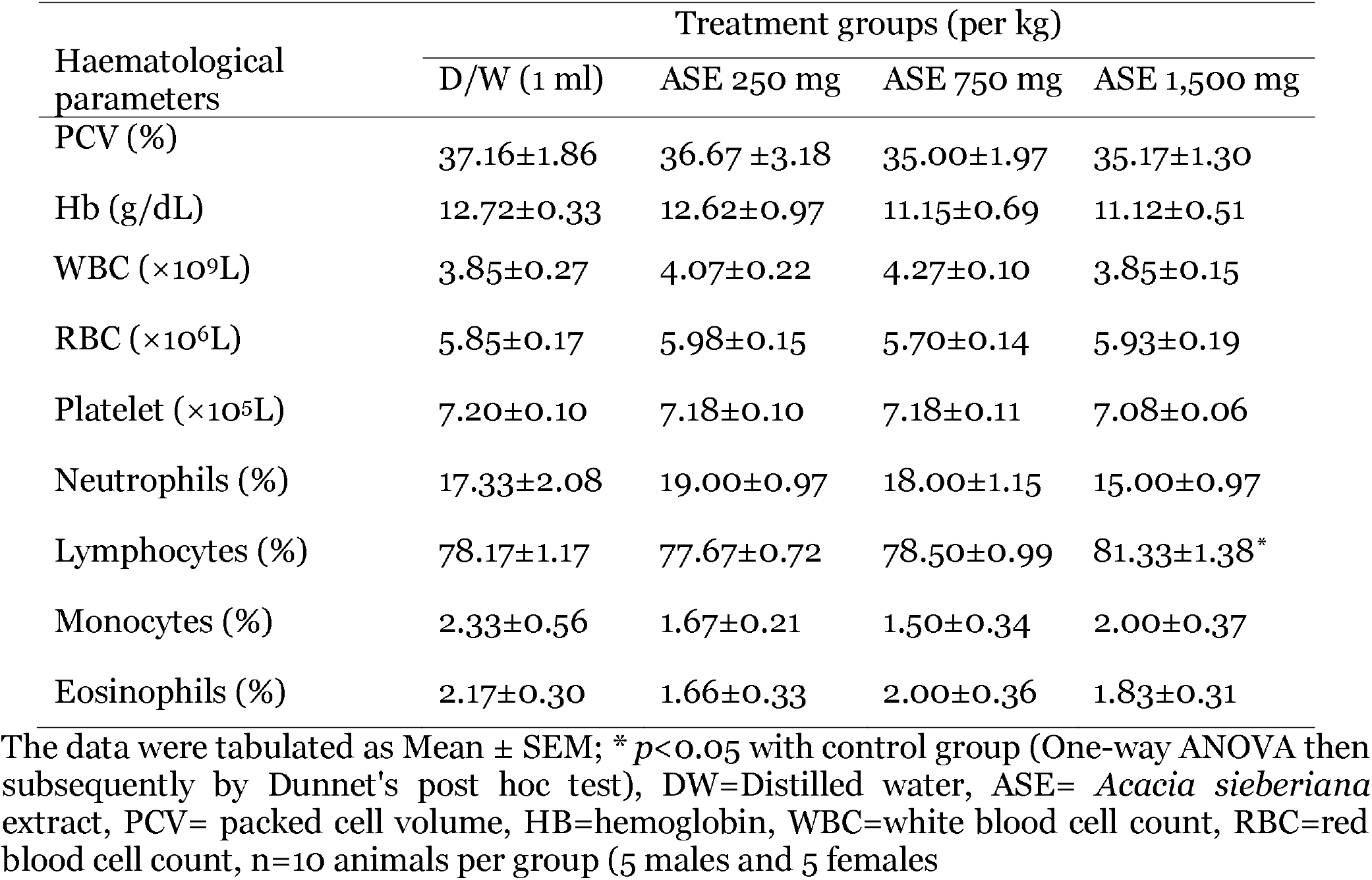
Haematological parameters.

| Haematological parameters | Treatment groups (per kg) |  |  |  |
| --- | --- | --- | --- | --- |
|  | D/W (1 ml) | ASE 250 mg | ASE 750 mg | ASE 1,500 mg |
| PCV (%) | 37.16±1.86 | 36.67 ±3.18 | 35.00±1.97 | 35.17±1.30 |
| Hb (g/dL) | 12.72±0.33 | 12.62±0.97 | 11.15±0.69 | 11.12±0.51 |
| WBC (×10<sup>9</sup>L) | 3.85±0.27 | 4.07±0.22 | 4.27±0.10 | 3.85±0.15 |
| RBC (×10<sup>6</sup>L) | 5.85±0.17 | 5.98±0.15 | 5.70±0.14 | 5.93±0.19 |
| Platelet (×10<sup>5</sup>L) | 7.20±0.10 | 7.18±0.10 | 7.18±0.11 | 7.08±0.06 |
| Neutrophils (%) | 17.33±2.08 | 19.00±0.97 | 18.00±1.15 | 15.00±0.97 |
| Lymphocytes (%) | 78.17±1.17 | 77.67±0.72 | 78.50±0.99 | 81.33±1.38* |
| Monocytes (%) | 2.33±0.56 | 1.67±0.21 | 1.50±0.34 | 2.00±0.37 |
| Eosinophils (%) | 2.17±0.30 | 1.66±0.33 | 2.00±0.36 | 1.83±0.31 |

The data were tabulated as Mean ± SEM; \* $p < 0.05$ with control group (One-way ANOVA then subsequently by Dunnet's post hoc test), DW=Distilled water, ASE= *Acacia sieberiana* extract, PCV= packed cell volume, HB=hemoglobin, WBC=white blood cell count, RBC=red blood cell count, n=10 animals per group (5 males and 5 females)

### 3.8 Histopathological findings on the liver

The liver photomicrograph of the animals revealed dose-dependent pathological alterations. The groups treated with the ASE revealed slight to moderate hepatocellular necrosis and slight vacuolation. The effects of the ASE on the hepatic histopathology following 28-days of repeated oral administration are presented in Figure 1.

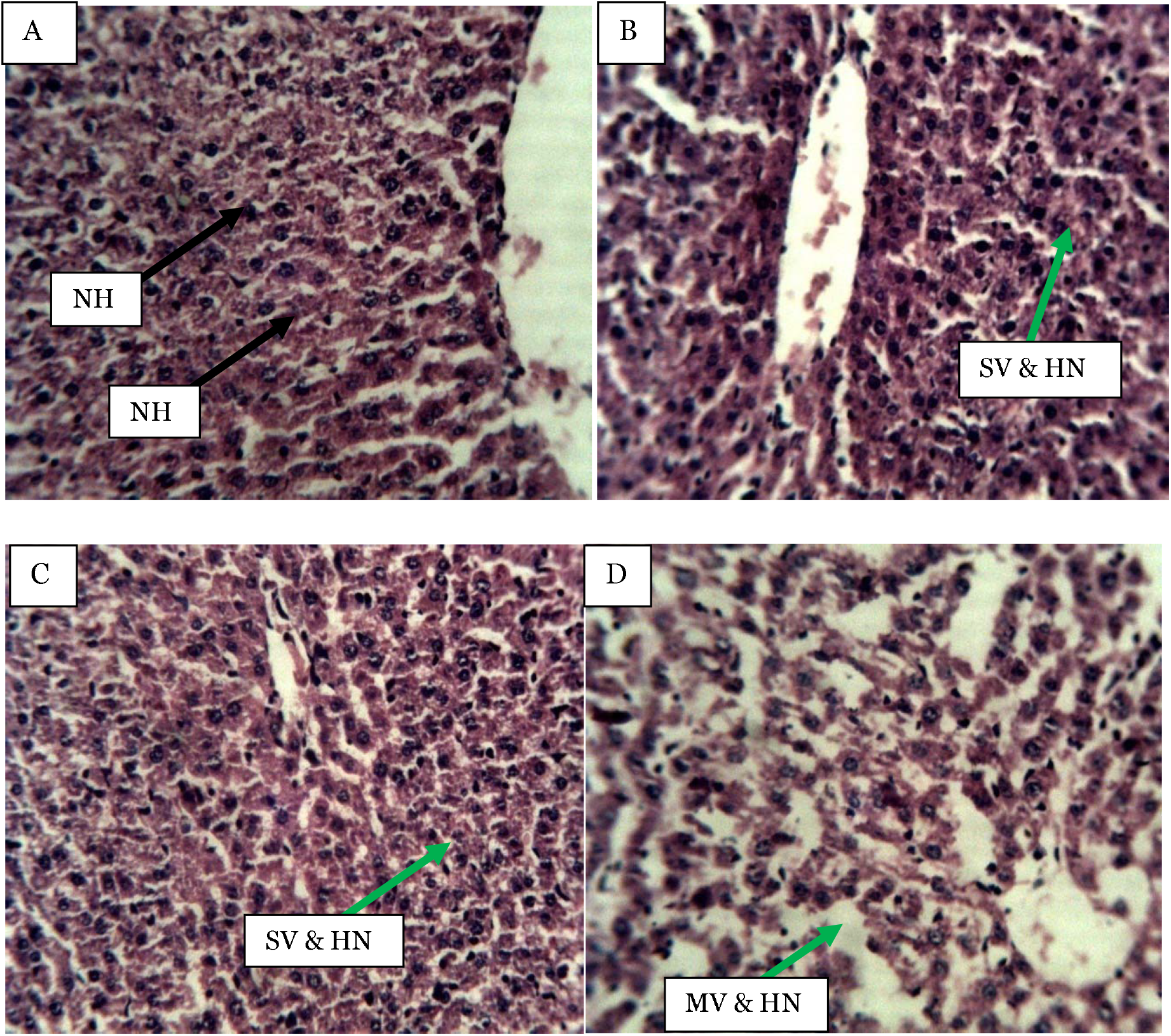

### 3.9 Histopathological findings on the kidney

The kidneys showed slight tubular distortion and necrosis as well as lymphocytes hyperplasia (250 mg/kg), slight glomerular necrosis (750 mg/kg), slight glomerular necrosis and lymphocyte hyperplasia (1,500 mg/kg). The effects of the ASE on the renal histopathology following 28-days of repeated oral administration are presented in Figure 2.

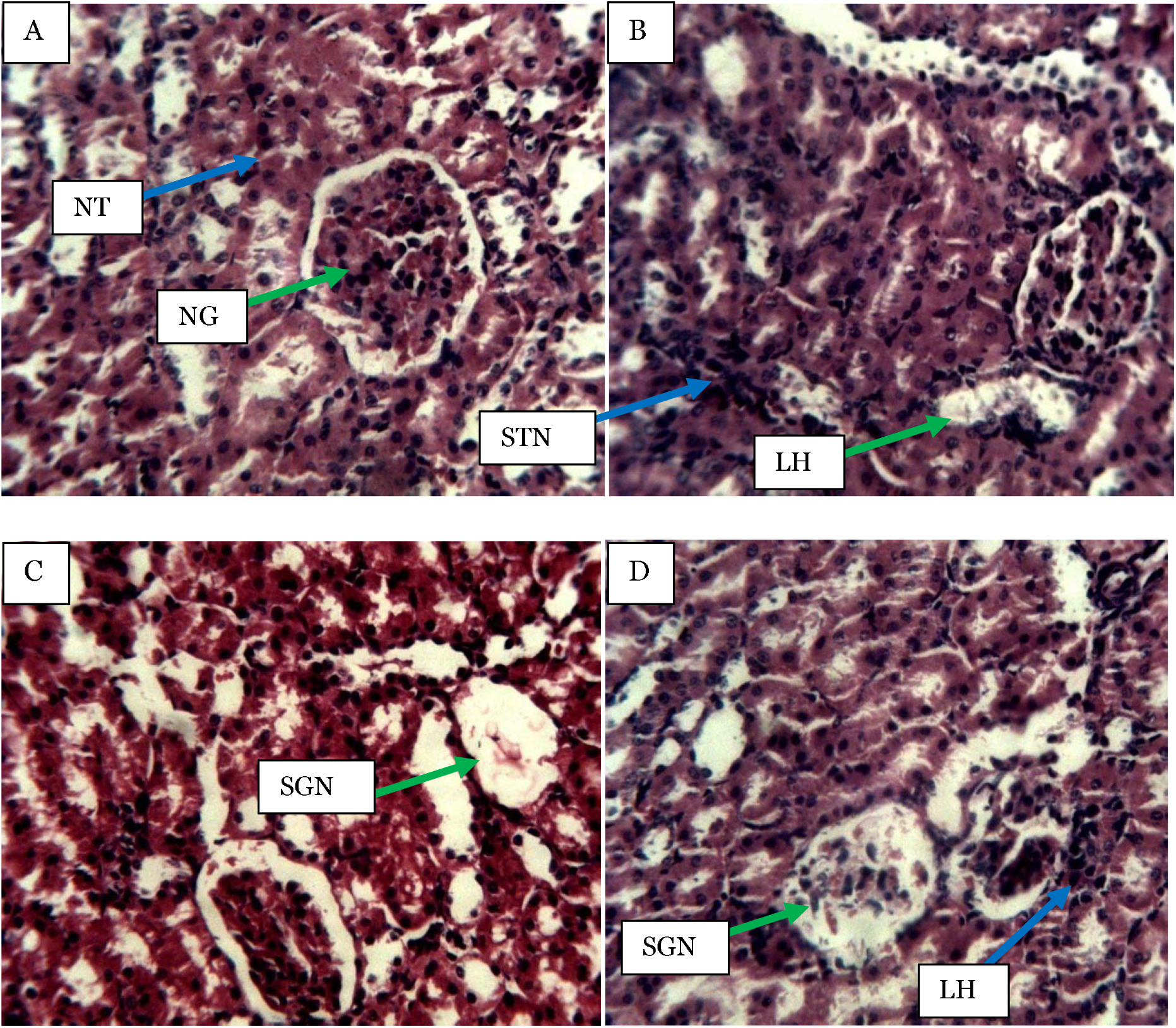

### 3.10 Histopathological findings on the lung

The lungs showed a dose-dependent pathological change (slight to moderate alveoli congestion and slight lymphocyte hyperplasia) in all the treated groups. The effects of the ASE on the lung histopathology following 28-days of repeated oral administration are presented in Figure 3.

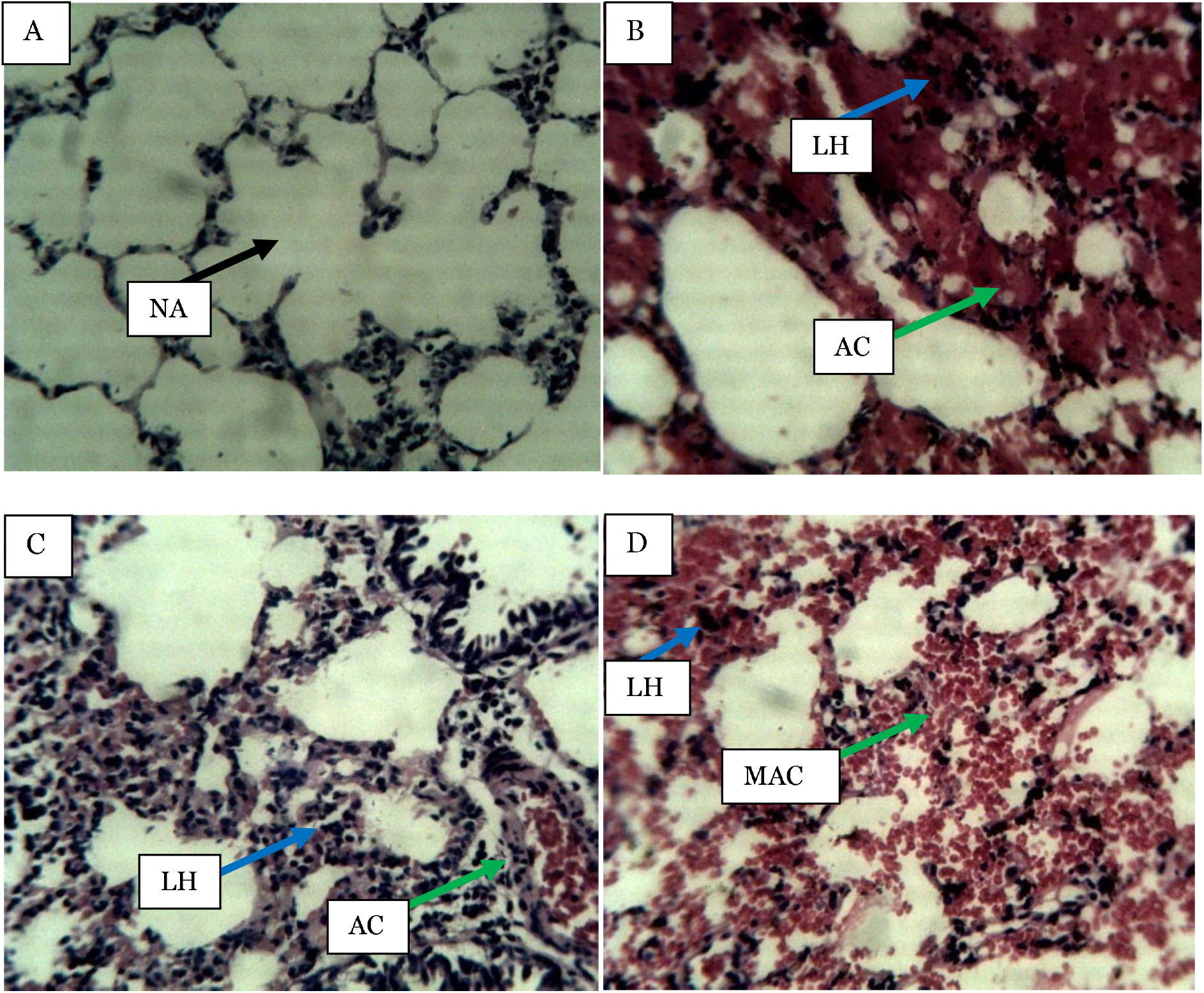

### 3.11 Histopathological findings on the heart

There were no histopathological alterations in the heart muscles of the animals in all the treated groups. The effects of the ASE on the histopathology of the heart muscles following 28-days repeated oral administration are presented in Figure 4.

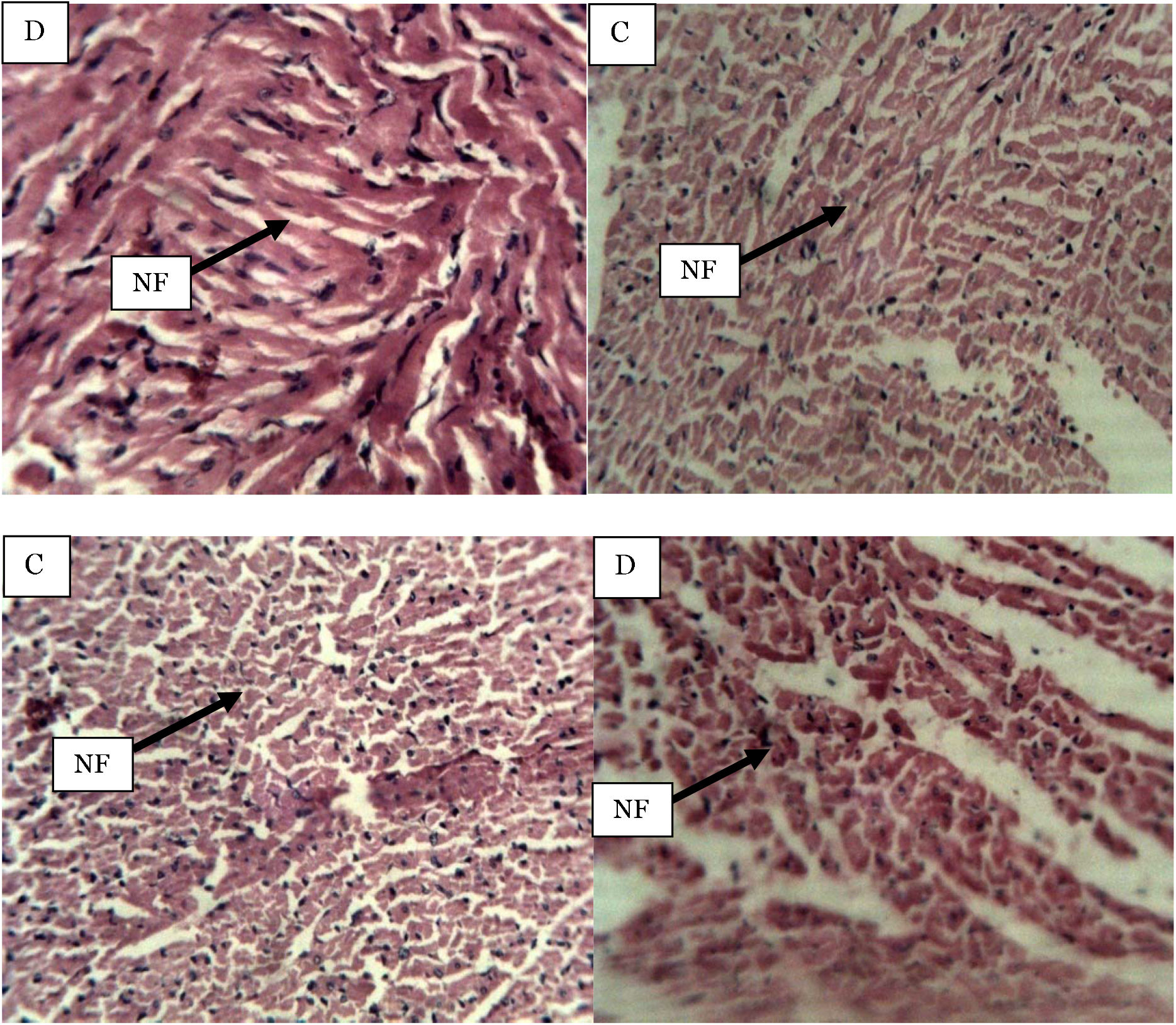

## 4.0 Discussion

The plants-sourced products possess therapeutic properties against many ailments (Jiménez-estrada et al., 2013; Miekus et al., 2020). However, research has shown some medicinal plants as toxic (Kharchoufa et al., 2018; Nasri & Shirzad, 2013; Ndhlala et al., 2013). Hence, there is a requirement to evaluate their potential harmful actions despite their medicinal properties (Ndhlala et al., 2013). There have been significant concerns about the use of medicinal plants due to a lack of evidence-based information, including regulatory and legal concerns, pharmacovigilance, and the paucity of information on their safety (Kale et al., 2019; Mazumder et al., 2016). The Food and Drug Administration (FDA) has advised appropriate precautionary measures for the unregulated utilization of herbal substances due to their probable toxicity (De Smet, 2004; Kale et al., 2019). In evaluating toxic potentials of herbal preparations, general behaviour, weekly body weight, biochemical and blood biomarkers together with histological results are useful (Jothy et al., 2011). As such, the current work intends to check the sub-acute oral toxic effects of *Acacia sieberiana*.

Natural products including plants, contain chemical agents commonly known as secondary metabolites that play a key role in drug development (Susanto et al., 2017). Identifying biological compounds in a plant is necessary for further pharmacological studies (Momin et al., 2014). The determination of phytochemical agents in the present research indicates that the ASE has cardiac glycosides, triterpenes, saponins, tannins, flavonoids and alkaloids. These phytocompounds could be responsible for the vast biological actions of plants (Kpemissi et al., 2020). Even though these chemical compounds could possess potential therapeutic effects, they may pose harmful risks to living organisms (Kpemissi et al., 2019). For example, cardiac glycosides possess heart-related toxicities (Botelho et al., 2018), tannins pose a severe risk to the liver and kidney (Ekambaram et al., 2018), whereas flavonoids are associated with liver diseases, anaemia and hypoglycaemia (Galati & Brien, 2004). Also, saponins and alkaloids produce liver disturbances (Qin et al., 2009; Wiedenfeld, 2011).

The acute toxicity investigation of therapeutic agents helps uncover possible harmful actions after a short term administration at a single dosage. Besides, it is employed in the first stage to investigate the pharmacological effects of new therapeutic compounds, particularly LD_50_ determination (Kpemissi et al., 2020; Musila et al., 2017; Ugwah-oguejiofor et al., 2019). Therefore, the non-toxic effects and lack of death by the ASE after the acute administration indicates that its LD_50_ may be more than 5,000 mg/kg. The findings concur with a previous report on *Piper capense* (Wamba et al., 2020), *Combretum hypopilinum* (Ahmad et al., 2020) and *Stachytarpheta cayennensis* (Olayode et al., 2020).

The subacute toxicity evaluation is used to uncover the possible harmful effects of chemical agents after repeated administration over a period of 28-days in rodents (Christapher et al., 2017). Besides, it gives information on the cumulative effects of the test agents and the specific organs (Loha et al., 2019). In conducting toxicological studies of extracts and drugs, evaluation of body weight, biochemical biomarkers such as liver and kidney functions as well as blood indices play an important role due to their response to the effects of toxic agents (Loha et al., 2019; Rahman et al., 2001). Based on the current work, no signs of harmful consequences of chemicals and mortality were seen over the 28-days of the experimental duration.

The change in the weekly body weight is an index for possible poisonous effects from chemical agents as a result of fat deposition, loss of appetite, and low caloric intake (Prasanth et al., 2015). Other secondary metabolites, including tannins and saponins, interfere with nutritional intake and could produce weight loss (Nguenang et al., 2020). Therefore, in the current work, the decline in the body weight produced by the ASE may be related to loss of appetite that declines caloric intake and impairs nutrient absorption. Other medicinal plants such as *Stachytarpheta cayennensis* (Olayode et al., 2020), *Epigynum auritum* (Yang et al., 2019), and *Bridelia ferruginea* (Bakoma et al., 2013) were documented to have reduced body weight.

The liver is among the essential body organs that play many biological activities such as biotransformation, secretion, storage and detoxification of unwanted substances (Khan et al., 2019). Some medicinal plants target the liver to manifest their toxic consequences (Quan et al., 2020). The plant-based bioactive agent-induced liver injury has been an important cause for the termination of the drug discovery process because these agents target important hepatic parameters (He et al., 2019; Tang et al., 2020). The ALT, AST, and ALP are liver biomarkers that determine the liver metabolic activities (El Kabbaoui et al., 2017) and are considered to investigate and manage liver disorders (Kim et al., 2012; Villela-nogueira et al., 2005). The plasma levels of these hepatic enzymes elevate as a result of cell membrane permeability alterations (Li et al., 2019). On the contrary, their decline shows chronic renal disorder, worsening liver disease (Cavalcanti et al., 2012). Besides, the ALP is vital to diagnose biliary duct diseases (El Kabbaoui et al., 2017; Ray et al., 2017). The moderate increase in ALP, with little or no increase in ALT and AST suggests primary biliary cirrhosis or primary sclerosing cholangitis (Hall & Cash, 2012). Therefore, the dose-dependent elevation of the ALP and the normal ALT and AST levels in the current work reveal that the ASE could cause biliary cirrhosis disease. The elevated ALP in this research is in line with a report on *Melastoma malabathricum* (Kamsani et al., 2019), *Dicoma anomala* (Balogun et al., 2016) and *Haloxylon scoparium* (Kharchoufa et al., 2018).

The renal system plays an essential function in regulating key physiological activities, including acid-base balance regulation and fluid and electrolytes maintenance (Bencheikh et al., 2021). The principal function of the kidney is to remove unwanted metabolic substances from the body system (Wamba et al., 2020). Chemicals produce renal toxic effects due to the incapability of the kidney to remove unwanted products (Kim & Moon, 2012). Hence, the evaluation of renal parameters provides data on the appropriate kidney functions (Wamba et al., 2020). The urea is produced in the liver from ammonia as a metabolite of protein biotransformation, which is excreted by the kidneys via glomerular filtration. It is used to diagnose renal disorders (Aprioku et al., 2014; Oyagbemi et al., 2013). In renal diseases, the urea level in the plasma is elevated (Oyagbemi et al., 2013). Therefore, the increase in plasma urea level at 1,500 mg/kg in this work indicates that the ASE may cause renal damage as a result of increased urea production beyond its excretion. The findings could be supported by renal tubular distortion and necrosis observed in the renal histopathology. Besides, the elevated plasma urea level was observed in some other medicinal plants such as *Simarouba glauca* (Osagie-eweka et al., 2021), *Terminalia schimperiana* (Awotunde et al., 2019) and *Caralluma dalzielii* (Ugwah-oguejiofor et al., 2019).

Haematological investigations are employed in determining the poisonous consequences of chemical substances on blood, such as plant-derived agents (Loha et al., 2019). A change in haematological indices could be associated with diseases that affect the hematopoietic system because of its sensitivity to harmful chemical substances including drugs (El Kabbaoui et al., 2017; Olorunnisola et al., 2012). Besides, the leucocytes including lymphocytes, protect the body from invading organisms, inflammation and tissue damage (Hervé et al., 2020). The current research indicates that the ASE may not lead to anaemia. The extract could have bioactive agents with immune system boosting effects which agrees with findings on *Lycopersicon esculentum* (Nguenang et al., 2020).

Histopathological analysis of essential body organs forms part of a crucial toxicological evaluation of biological compounds (Traesel et al., 2016). Hepatic necrosis results from the hepatic inflammatory reaction, mononuclear and neutrophils recruitment in the liver (Sharifudin et al., 2013). The hepatic necrosis seen in the current study supports the potentially harmful actions of the ASE on the liver tissues. The hepatocellular necrosis and vacuolations in the liver may have resulted in increased ALP levels. It had been reported that liver ALP is located in the microvilli of bile canaliculi and on the sinusoids surface of the hepatocytes (Hall & Cash, 2012); thus agents that affect the liver as with ASE may also cause alterations in the concentration of this enzyme. The results indicate the potential harmful effects of the ASE on the liver which corroborates with the elevated ALP levels.

The slight tubular and glomerular necrosis seen in the current research could be due to the poisonous compounds in the ASE reaching the kidney via the systemic circulation that may cause dysfunction of kidney tubules (Zhao et al., 2007). The tubular and glomerular damage may interfere with electrolytes reabsorption and glomerular filtration, and could be accompanied by the elevated urea level observed (Zhao et al., 2007). The alveoli congestion in the lungs produced by the ASE could have prevented adequate oxygen and other gases from being absorbed into the pulmonary circulation for normal body functions. Besides, the lack of histopathological effects of the ASE on the heart muscle indicates an absence of cardiotoxicity as a result of lack impairment with erythropoiesis related to the normal delivery of blood to the myocardium.

## 5.0 Conclusions

The findings in this work have revealed that ASE may be safe after acute exposure. On the contrary, the extract could be harmful on sub-acute administration, particularly to the liver and kidney. Hence, further investigations should be conducted to check the chronic toxicity of the plant. In addition, the traditional herbalist should be well informed on the potential toxic consequences of the plant for long term use.

## List of abbreviations

ABU: Ahmadu Bello University
ABUCAUC: ABU Ethical Committee on Animal Use and Care Research Policy
ALP: Alkaline phosphatase
ALT: Alanine transaminase
ALT: Aspartate transaminase
ANOVA: One-way analysis of variance
ARRIVE: Animal Research: Reporting of *In Vivo* Experiments
ASE: *Acacia sieberiana* extract
D/W: Distilled water
EDTA: Ethylenediaminetetraacetic acid
H&E: Haematoxylin and eosin
HB: Hemoglobin
LD50: Median lethal dose
OECD: Organization of Economic Co-operation and Development
RBC: Red blood cell
SEM: Standard error of mean
WBC: White blood cell

## Declaration

## Ethics approval and consent to participate

The permission for the experiment was given by the Ahmadu Bello University Ethical Committee on Animal Use and Care Research Policy (approval number: ABUCAUC/2016/049) and carried out as per the Animal Research: Reporting of I*n Vivo* Experiments (ARRIVE) protocols.

## Consent for publication

Not applicable

## Availability of data and material

The datasets generated during and/or analyzed during the current study are available from the corresponding author on reasonable request

## Authors’ Contribution

**MW:** Conceptualization, investigation, resources, data curation, writing and data analysis. **JIE:** Validation, supervision, project administration, and review. **AM:** Investigation and review. **MHA:** Writing of the original draft, critically revised the whole manuscript and editing. All the authors read and approved the final manuscript

## Acknowledgements

The authors especially thank all staff of the Pharmacology and Therapeutics Department, ABU, Nigeria, for the support throughout the research period.

